# Gut microbiota requires vagus nerve integrity to promote depression

**DOI:** 10.1101/547778

**Authors:** Eleni Siopi, Soham Saha, Carine Moigneu, Mathilde Bigot, Pierre-Marie Lledo

## Abstract

Recent studies have shown that gut microbiota (GM) can influence hippocampal plasticity and depression-like behaviors (1,2,3), yet the underlying mechanisms remain elusive. The gastrointestinal branches of the vagus nerve, which constitute a direct bidirectional route of communication between the gut and the brain, are well suited to carry neural messages associated with changes in peripheral states to the brain (4,5). For instance, the ability of some gut bacterial strains to induce anxiety-like behaviors and alter the expression of proteins in the hippocampus depends on vagal afferents (6,7). Similarly, gut vagal sensory signaling affects adult hippocampal plasticity and neurogenesis (8). Impairment in brain plasticity is known to generate depressive states in mice (9,10). Yet, it is not known whether GM abnormalities require a functional, intact vagus nerve to promote depression or if GM borrows other pathways, such as humoral immune responses. Here we used the fecal transplantation paradigm to study whether changes in GM necessitate the integrity of the vagus nerve to impact on adult hippocampal neurogenesis and depressive-like behavior. Alteration of GM was induced by unpredictable chronic mild stress (UCMS), a standard model of depression (11).

We used mice (8 weeks old, C56BL6/j) that sustained 9 weeks of UCMS (UCMS group), and their controls (CT group), collectively called “microbiota donor” mice. Fresh fecal samples were harvested from microbiota donor mice at the end of the 9^th^ week of UCMS and were transferred by oral gavage to “microbiota recipient” mice (8 weeks old, C56BL6/j), which had been treated with broad-spectrum antibiotics (abx) (12) during one week prior the fecal transfer. Abx was discontinued 24h prior microbiota transplantation. Microbiota recipient mice received either the CT (CT-tr group) or the UCMS (UCMS-tr group) fecal suspension at 1 and 4 days following abx discontinuation (Figure 1A,B). In order to assess whether the microbiota transfer can affect behavior and neurogenesis via the vagus nerve, we distinguished two different subgroups of microbiota recipient mice: animals that had sustained subdiaphragmatic vagotomy (Vx) 4 weeks prior fecal transplantation, and a sham group (Figure 1A,B). Microbiota recipient mice were maintained in isolators for 7 weeks following the fecal transfer.

**Figure 1.**
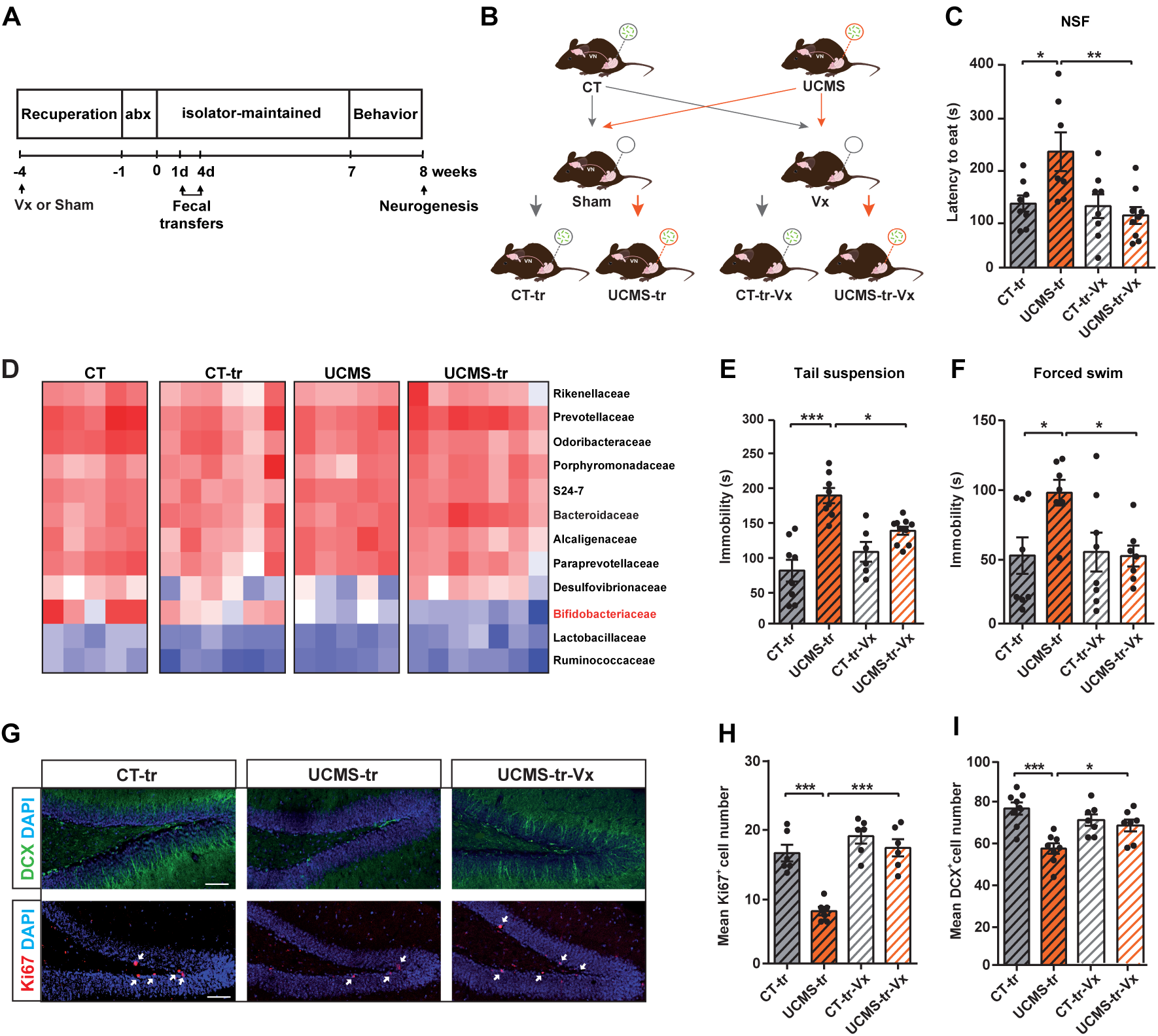
Gut microbiota from chronically stressed mice generates depression and decreases hippocampal adult neurogenesis through the vagus nerve. **(A)** Experimental design used for assessing the microbiota changes, behavioral responses and hippocampal neurogenesis in mice that were colonized by microbiota from either CT or UCMS mice. Microbiota recipient mice were treated with broad spectrum antibiotics (abx) for one week prior fecal transfer and were kept in isolators in order to maintain their bacterial profile. Mice sustained Vx at 4 weeks prior fecal inoculation. **(B)** Schematic representation of the fecal transplantation paradigm and the different groups used in the study. **(C)** In the NSF test, mice that were inoculated with UCMS-derived microbiota displayed an increased latency to eat the pellet (\**P*<0.05) that was completely abolished by Vx (\*\**P*<0.01). **(D)** Heatmap showing the mean relative abundances of the different bacterial families in the 16S rRNA sequencing across all the experimental replicates. **(E, F)** In the tail suspension and forced swim tests, which are both indicative of despair-like behavior, UCMS-tr mice displayed increased immobility (\*\*\**P*<0.001 and \**P*<0.05), and Vx abolished this behavior (\**P*<0.05). **(G)** Representative images of Ki67^+^ and DCX^+^ cells in the dentate gyrus of CT-tr, UCMS-tr and UCMS-tr-Vx mice. Scale bars: 100 mm. **(H)** UCMS-microbiota transfer decreased the number of Ki67^+^ proliferating cells in the dentate gyrus of recipient mice (\*\*\**P*<0.001) but not in mice that sustained Vx. **(I)** Transfer of UCMS microbiota decreased the number of DCX^+^ immature neurons in the dentate gyrus (\*\*\**P*<0.001). Vx abolished this decrease (\**P*<0.05). All data are represented as mean ± SEM. Statistical significance was calculated using two-way ANOVA followed by Tukey post-hoc test. abx: antibiotics, NSF: novelty suppressed feeding, Vx: vagotomy.

To assess the impact of fecal transfer on GM composition, we performed 16S rRNA gene V3-V4 region sequencing on the fecal samples. Taxonomic analysis of bacterial families revealed differences between CT and UCMS GM compositions, which were transferred to their respective recipient mice, with the most significant change being observed in Bifidobacteriaceae (Figure 1D). More precisely, both UCMS and UCMS-tr mice had significantly lower levels of Bifidobacteriaceae compared to CT and CT-tr mice respectively (*P*=0,0016 *and P*=0,0039, Tukey post-hoc). This finding supports previous studies showing that Bifidobacteria and prebiotics exert anti-depressant effects in rats (13, 14).

We next sought to determine whether these UCMS-induced GM changes are sufficient to change behavior. We employed a battery of standard behavioral tests to assess anxiety- and depression-like behaviors. Our results showed that UCMS microbiota recipient mice adopted the behavioral phenotype of donor mice, characterized by depressive-like responses, and that transmission of the depressive-like phenotype was abrogated by Vx (Figure 1C,E,F). For instance, in the novelty suppressed feeding (NSF) test (Two-way ANOVA, F(1, 28)=6.5; *P*=0.02), Tukey's post-hoc showed that while the transfer of UCMS microbiota significantly increased the latency to eat (*P*<0.05), Vx completely reversed this phenotype (*P*=0.005) (Figure 1C). Moreover, in both the tail suspension and forced swim tests [Two-way ANOVA, F(1, 28)=10.99, *P*=0.003; and F(1, 26)=4.23; *P*=0.05 respectively], UCMS-tr mice were more immobile compared to their control counterparts (*P*<0.001; *P*=0.02), and Vx abolished the phenotype (*P*=0.02; *P*=0.03) (Figure 1E,F). Taken together, our observations indicate that UCMS-induced changes in GM lead to depressive states only if the vagus nerve remains functional.

To further investigate whether these depressive states are accompanied by changes in adult hippocampal neurogenesis, we performed immunostaining for Ki67, a marker of proliferating cells, and doublecortin (DCX), a marker of transient proliferating neuronal progenitor cells, in the dentate gyrus (DG) of the hippocampus. The large pool of DCX^+^ cells in the DG, besides being a step in the maturation process of new neurons, is of importance in buffering the negative feedback on the hypothalamic–pituitary-adrenal axis (15). Counting of Ki67^+^ cells in the DG [Two way ANOVA, F(1, 24)=9.96; *P*=0.005], revealed that the transfer of UCMS-derived microbiota significantly decreased Ki67^+^ cell number (*P*<0.001), and that this decrease was completely abolished by Vx (*P*<0.001) (Figure 1G,H). Our results on the number of DCX^+^ cells went on the same line. UCMS-tr mice had a significant decrease in DCX^+^ cells (*P*=0.001) that was attenuated by Vx (*P*=0.03) (Figure 1G,I). These data show that vagal nerve activity is essential to mediate the impact of gut dysbiosis on hippocampal adult neurogenesis.

The present study demonstrates that chronic stress induces changes in GM composition, notably diminution of the Bifidobacteriaceae family. When transfered to healthy recipient mice, these changes promote depressive states and decrease adult hippocampal neurogenesis. While the exact underlying cascades of events are not elucidated, we found that the transmitted dysbiosis requires an intact vagus nerve to be effective. Our results are in line with recent studies showing altered cecal and fecal microbiota composition in experimental models of stress (16,17) and in depressed patients (18). These alterations may contribute to the neuroprogression of stress-related depression by altering physiological and/or metabolism processes that activate vagal afferents, such as microbial peptides, neurotransmitter release and immunity/inflammation (19,20). Our results are in line with recent data showing that the vagus nerve mediates the anxiolytic effects of some probiotic strains in rodents (6,21,22). The two existing studies pertaining to the effect of the vagus nerve on hippocampal plasticity show contradictory results, (23, 24). While the reasons underlying these discrepancies are not clear, they may be due to methodological differences, such as the Vx procedure itself. Relevant electrophysiological data on vagal tone should be implemented in future studies to decipher how GM promote depression. Our data further support the emerging hypothesis that GM can influence brain function and behavior directly through a nervous pathway mediated by vagal afferents.

## Acknowledgements and disclosures

We thank all of Pierre-Marie Lledo’s lab members for helpful discussions during the course of this study. We also want to thank the members of the Institut Pasteur Animal Facility who were essential for this project, and in particular Marion Bérard, Martine Jacob, Thierry Angelique and Eddie Maranghi. This work was supported by the *Investissements d'Avenir* program managed by the *Agence Nationale de la Recherche* (ANR) under the reference ANR-11-IDEX-0004-02 and ANR-10-LABX-73, the *Agence Nationale de la Recherche* (ANR-15-CE37-0004-01) and the Life Insurance Company “AG2R-La Mondiale”.

The authors declare to have no biomedical financial interests or potential conflicts of interest.

## Author contributions

ES initiated the project in the P-M.L. laboratory, designed the study, performed the experiments and wrote the manuscript. S.S. analyzed the 16S data. C.M. contributed in the subdiaphragmatic vagotomy surgeries. M.B. contributed in some experiments. P-M.L. supervised the study and edited the manuscript with input from the other authors.

